# Mapping the Subtype-Specific PARP1 ADP-ribosylated Proteome in Breast Cancer Cells

**DOI:** 10.1101/2025.09.30.679484

**Authors:** Sneh Koul, Minjung Kwon, Poulami Tapadar, Yangyang Dai, Tulip Nandu, Dan Huang, Cristel V. Camacho, W. Lee Kraus

## Abstract

Breast cancers are molecularly heterogeneous, with subtype-specific differences in transcriptional programs, chromatin architecture, and therapeutic responses. While PARP1 has been extensively studied in the context of DNA repair, emerging evidence implicates its catalytic activity in a broader set of cellular processes, including the regulation of gene expression. Here, we employed an NAD analog-sensitive PARP1 (asPARP1) chemical genetics approach combined with mass spectrometry to map the ADP-ribosylated proteome across six human breast cancer cell lines representing luminal and basal/triple negative subtypes. We identified thousands of PARP1 substrates and hundreds of Glu/Asp ADPRylation sites, revealing both shared and subtype-specific modifications in cell lines maintained under basal growth conditions. Luminal-specific substrates were enriched in chromatin and transcriptional regulators, whereas basal-specific substrates were preferentially linked to translation and RNA processing, highlighting lineage-dependent PARP1 activity. Transcription factors emerged as major substrates, with TFAP2A serving as a proof-of-concept; it is selectively ADPRylated in luminal cells and inhibition of PARP1-mediated ADPRylation modulates its promoter occupancy in a subtype-specific manner. Our data provide a new resource for studying subtype-specific PARP1-mediated ADPRylation in breast cancer cells. Collectively, our findings expand the conceptual framework for PARP1 function beyond DNA repair, offering mechanistic insights into subtype-specific gene regulation and potential determinants of PARP inhibitor sensitivity in breast cancer.

## Introduction

The molecular landscapes of cells within a particular cancer govern pathogenesis and response to drug treatments. Gene expression profiling analyses have been used to characterize the molecular heterogeneity in cancers, including multiple molecular subtypes of breast cancer, which have distinct hormonal and therapeutic responses, patient prognoses, and patterns of gene expression (1,2). The luminal breast cancer subtypes, which express hormone receptors (e.g., estrogen receptor alpha and progesterone receptor; ERα and PR, respectively), are the most commonly diagnosed subtypes (2,3). In contrast, the basal/triple negative breast cancer subtypes are hormone receptor-negative and associated with a much poorer prognosis (4–6). In addition, they are more metastatic, with the lowest survival rates and a propensity for acquiring resistance to drug treatments (4,6,7). In this study, we explore the possibility that profiling post-translational modifications (PTMs) of proteins, such as ADP-ribosylation (ADPRylation), can provide an alternate way to characterize the molecular heterogeneity of breast cancers.

ADPRylation is a reversible PTM of substrate proteins resulting in the covalent attachment of a single ADP-ribose (ADPR) (monoADPR or MAR) or polymers of ADPR units (polyADPR or PAR) derived from nicotinamide adenine dinucleotide (NAD^+^) on a variety of amino acid residues (8,9). ADPRylation is catalyzed by the Poly(ADP-ribose) Polymerase (PARP) family of enzymes, consisting of 17 members that have distinct structural domains, activities, subcellular localizations, and functions (8,10–12). Early studies into PARP functions focused primarily on the biochemistry and molecular biology of PARP1 (the founding member of the family) in DNA repair (13). In this context, inhibition of PARP1 can selectively induce synthetic lethality in cancer cells that harbor homologous recombination deficiencies (HRD), such as mutations in *BRCA1/2* genes (14,15). This has led to the FDA approval of 4 different PARP inhibitors in the last 10 years, which have shown promise in the treatment of ovarian and breast cancers, as well as other cancer types (16–18).

However, a growing number of studies have demonstrated efficacy of PARP inhibitors irrespective of *BRCA1/2* or HRD status: Olaparib, rucaparib and niraparib in recurrent/relapsed ovarian cancer (19–23), and niraparib in newly diagnosed advanced ovarian cancer (24), for example, suggesting therapeutic potential of PARP inhibitors beyond DNA repair mechanisms (25,26). In this regard, the mechanistic and biological understanding of the role of PARPs in a wide range of biological processes has grown considerably beyond DNA repair (11,26,27), and these roles also impact responses to PARP inhibitors (28). A number of seminal studies on PARP1 over the past decade have shifted the focus from DNA repair to the regulation of chromatin structure and gene regulation (26,29,30). In breast cancer, we have previously shown that PARP inhibitors can attenuate estrogen-dependent growth of multiple ERα-expressing (ER+) breast cancer cell lines that lack known *BRCA1/2* mutations and are proficient in HR (31). Many of the gene regulatory functions of the nuclear PARPs requires their catalytic activity (9,26,30,32).

Given that the molecular landscapes of cells within a particular cancer govern pathogenesis and response to drug treatment, our goal is to identify breast cancer subtype-specific gene regulatory substrates that are ADPRylated by PARP1 and may influence response to PARP inhibitors. We have used an NAD^+^ analog-sensitive PARP1 (asPARP1)-mass spectrometry approach that we developed previously (9,33) to map the ADPRylated proteome across six human breast cancer cell lines representing luminal (MCF-7, T-47D, ZR-75) and basal/triple-negative (HCC70, MDA-MB-231, MDA-MB-468) breast cancer subtypes. Our study contributes a rich proteomic resource that can provide a more functional understanding of the effects of PARP1 substrate ADPRylation on the response of breast cancer cells to clinically used PARP inhibitors.

## Materials and Methods

### Antibodies, drugs, and chemicals

The antibodies used for Western blotting were anti-PARP1, purchased from Active Motif (39559; RRID:AB_2793257), the custom recombinant antibody-like anti-poly-ADP-ribose binding reagent (anti-PAR) was generated and purified inhouse (now available from EMD Millipore, MABE1031; RRID:AB_2665467), β-tubulin (Abcam, ab6046; RRID:AB_2210370), TFAP2A (Proteintech, 13019-3-AP; RRID:AB_2199414), TFAP2A conjugated to agarose (Santa Cruz, sc-12726 AC; RRID:AB_667767), and β-actin (Abcam, ab8226; RRID:AB_306371). Secondary antibodies for Western blotting included goat anti-rabbit HRP-conjugated IgG (Pierce, 31460; RRID:AB_228341), goat anti-mouse HRP-conjugated IgG (Pierce, 31430; RRID:AB_228307). For Western blotting, the primary antibodies were used at a 1:3000 dilution in 3% non-fat milk made in TBS-T, with subsequent detection using an appropriate HRP-conjugated secondary antibody used at a 1:6000 dilution in 3% non-fat milk made in TBST. Cells were treated with DMSO (vehicle), PARP inhibitors Olaparib (MedChemExpress; HY-10162) or Talazoparib (MedChemExpress; HY-16106) at indicated times and doses, and 1mM H_2_O_2_ for 5 min. The unnatural NAD^+^ analog 8-Bu(3-yne)T-NAD^+^ was purchased from the BIOLOG Life Science Institute (LSI), Bremen, Germany.

### Cell culture

Cell lines MCF-7 (HTB-22; RRID:CVCL_0031), T-47D (HTB-133; RRID:CVCL_0553), ZR-75-1 (CRL-1500; RRID:CVCL_0558), HCC70 (CRL-2315; RRID:CVCL_1270), MDA-MB-231 (CRM-HTB-26; RRID:CVCL_0062), MDA-MB-468 (HTB-132; RRID:CVCL_0419), and HEK-293T (CRL-3216; RRID:CVCL_0063) cell lines were purchased from the American Type Culture Collection and used for nuclear extract preparation, western blots, proliferation assay, and the variety of assays described in methods. MCF-7 cells were maintained in Minimum Essential Medium Eagle (Sigma-Aldrich, M1018) supplemented with 5% calf serum (Sigma-Aldrich, C8056). T-47D, ZR-75-1, HCC70 and HEK-293T were maintained in RPMI media (Sigma-Aldrich, R8758) supplemented with 2 mM Glutamax, 10% fetal bovine serum (FBS; Sigma-Aldrich, F0926) and 1% penicillin/streptomycin (P/S). MDA-MB-231 cells were maintained in DMEM (Sigma-Aldrich, D5796) supplemented with 10% FBS, 2 mM Glutamax, 1% P/S and 25 μg/mL gentamicin. MDA-MB-468 cells were maintained in alpha-MEM (Sigma-Aldrich, M8042) supplemented with 10% FBS, 0.1 mM HEPES, 1% non-essential amino acids, 2 mM glutamine, 1% sodium pyruvate, 1 μg/mL insulin, 1 ng/mL hydrocortisone, 12.5 ng/mL epidermal growth factor and 1% P/S. Fresh cell stocks were regularly replenished from the original stocks. All cell lines were verified for cell type identity using the GenePrint 24 system (Promega, B1870), and confirmed as *Mycoplasma*-free every 6 months using the Universal Mycoplasma Detection Kit (ATCC, 30-1012K).

### Preparation of whole cell lysates

The collected cells were washed with ice-cold PBS and resuspended in Whole Cell Lysis Buffer [50 mM Tris-HCl pH 7.9, 500 mM NaCl, 1 mM CaCl_2_, 0.2% Triton X-100, 1X complete protease inhibitor cocktail (Roche, 11697498001), 250 nM ADP-HPD (Sigma, A0627, a PARG inhibitor), 10 mM PJ34 (Enzo Life Sciences, ALX-270-289, a PARP inhibitor)] and incubated for 15 min at room temperature with gentle mixing to lyse the cells and extract the proteins. The lysates were clarified by centrifugation in a microcentrifuge for 5 min at 4°C at full speed (∼ 12,000 x *g*).

### Preparation of nuclear and cytosolic extracts

The collected cells were washed with ice-cold PBS and resuspended in Isotonic Buffer (10 mM Tris-HCl pH 7.5, 2 mM MgCl_2_, 3 mM CaCl_2_, 0.3 M sucrose, with freshly added 1 mM DTT, 1X protease inhibitor cocktail, 250 nM ADP-HPD, and 10 mM PJ34), incubated on ice for 15 min, and lysed by the addition of 0.6% IGEPAL CA-630 with gentle vortexing. After centrifugation, the nuclei were collected by centrifugation in a microcentrifuge for 1 min at 4°C at 1,000 rpm. The supernatant was collected as the cytoplasmic fraction. The pelleted nuclei were resuspended in Nuclear Extraction Buffer (20 mM HEPES pH 7.9, 1.5 mM MgCl_2_, 0.5 M NaCl, 0.2 mM EDTA, 25% v/v glycerol, with freshly added 1 mM DTT, 1X protease inhibitor cocktail, 250 nM ADP-HPD, and 10 mM PJ34) and incubated on ice for 30 min. The chromatin and soluble nuclear fractions were separated by centrifugation (12,000 x g, 30 min, 4°C) and the soluble chromatin fraction was collected to store at -80°C after flash-frozen.

### Dose response and cell proliferation assays

For dose response assays, cells were seeded at low density. Increasing doses of inhibitors (Olaparib or Talazoparib) were applied and media with fresh drug was replaced every 2 days. Cells were fixed for 10 minutes with 4% paraformaldehyde at room temperature on Day 6 and stained with 0.1% crystal violet in 20% methanol solution. After washing with PBS to remove unincorporated stain, the crystal violet was extracted using 10% glacial acetic acid and the absorbance was read at 595 nm. All proliferation assays were performed a minimum of three times using independent plating to ensure reproducibility. Growth curves were plotted and IC50s were calculated using GraphPad Prism.

### Purification of PARP1 and asPARP1 expressed in Sf9 insect cells

Sf9 insect cells, cultured in SF-II 900 medium (Invitrogen), were transfected with 1 μg of bacmid driving expression of wild-type PARP1 and analog-sensitive PARP1 (asPARP1) using Cellfectin transfection reagent as described by manufacturer (Invitrogen). After three days, the medium was collected as a baculovirus stock. After multiple rounds of amplification of the stock, the resulting high titer baculovirus was used to infect fresh Sf9 cells to induce expression of PARP1 protein for three days. The PARP1 expressing Sf9 cells were then collected by centrifugation, flash-frozen in liquid N_2_, and stored at -80°C. PARP1 pellets were thawed on wet ice. The cells were resuspended in Lysis Buffer [20 mM HEPES, pH 7.9, 0.5 M NaCl, 4 mM MgCl_2_, 0.4 mM EDTA, 20% glycerol, 250 mM nicotinamide, 2 mM β-mercaptoethanol, 2X protease inhibitor cocktail (Roche)] and lysed by Dounce homogenization (Wheaton). The lysate was clarified by centrifugation, mixed with an equal volume of Dilution Buffer (20 mM HEPES, pH 7.9, 10% glycerol, 0.02% NP-40), sonicated, and then clarified by centrifugation again. The clarified lysate was mixed with anti-FLAG M2 agarose resin (Sigma), washed twice with FLAG Wash Buffer #1 (20 mM HEPES, pH 7.9, 150 mM NaCl, 2 mM MgCl_2_, 0.2 mM EDTA, 15 % glycerol, 0.01% NP-40, 100 mM nicotinamide, 0.2 mM β-mercaptoethanol, 1 mM PMSF, 1 μM aprotinin, 100 μM leupeptin), twice with FLAG Wash Buffer #2 (20 mM HEPES, pH 7.9, 1 M NaCl, 2 mM MgCl_2_, 0.2 mM EDTA, 15% glycerol, 0.01% NP-40, 100 mM nicotinamide, 0.2 mM β-mercaptoethanol, 1 mM PMSF, 1 μM aprotinin, 100 μM leupeptin), and twice with FLAG Wash Buffer #3 (20 mM HEPES, pH 7.9, 150 mM NaCl, 2 mM MgCl_2_, 0.2 mM EDTA, 15% glycerol, 0.01% NP-40, 0.2 mM β-mercaptoethanol, 1 mM PMSF). The FLAG-tagged PARP proteins were eluted from the anti-FLAG M2 agarose resin (Sigma, A2220) with FLAG Wash Buffer #3 containing 0.2 mg/mL FLAG peptide (Sigma, F4799). The eluted proteins (∼0.5 mg/mL) were distributed into single use aliquots, flash frozen in liquid N_2_, and stored at -80°C until use.

### In vitro PARP1 automodification reactions

One hundred nanograms of purified recombinant PARP1 protein (wild-type or analog-sensitive) was incubated in Automodification Buffer [30 mM HEPES, pH 8.0, 5 mM MgCl_2_, 5 mM CaCl_2_, 0.01% NP-40, 1 mM DTT, 100 ng/μL sonicated salmon sperm DNA (Stratagene), 100 ng/μL BSA (Sigma)] with 250 μM NAD^+^ or NAD^+^ analog at 25°C for 15 min. Automodification (ADPRylation of PARP1) was monitored by Western blot or in-gel fluorescence.

### LC-MS/MS for the identification of PARP1 substrates using an NAD^+^ analog-sensitive PARP1 (asPARP1) approach

A detailed protocol for the analog-sensitive PARP1 (asPARP) approach using 8-Bu(3-yne)T-NAD^+^ coupled with protein mass spectrometry was described previously (33,34). This method was used to identify substrates of PARP1 across six different breast cancer cells, as well as the specific amino acid residues modified by PARP1 in those substrates. First, the system using catalytic activity of asPARP1 and click chemistry were verified by in vitro automodification assay and in-gel fluorescence assay. After proof-of-method, nuclear extracts from individual cell line were subjected to asPARP1-mediated catalytic assays, followed by mass spectrometry to identify the ADPRylated proteins (trypsin digestion elution) and the specific sites of modification (hydroxylamine elution) as described previously (33,34). Peptides identified from samples prepared with wild-type PARP1 (Wt) were treated as non-specific background.

### Analysis of LC-MS/MS data

#### LC-MS/MS peptide and site identification

The sites of ADPRylation were obtained from LC-MS/MS analysis as described previously (35). Briefly, proteins from affinity-resin pull downs were filtered using an FDR cutoff of < 1%. Within each biological replicate, specifically bound proteins were defined using a threshold of log_2_(experimental/control) > 1. For each cell line, replicate-level enriched sets were intersected to derive a high-confidence consensus substrate list. Additionally, ADPRylated sites identified with a high FDR confidence rate (<1%) were retained for further analysis. All ADPRylation sites identified from both replicates were used in the data analysis. The software, scripts, and other information about the analyses can be obtained by contacting the corresponding author (W.L.K.).

#### Overlap of core and broadly conserved ADPRylated proteins in luminal and basal breast cancer cell lines

ADPRylated proteins that were found to be common amongst luminal and basal cell lines were overlapped and showcased in a Venn diagram generated with bioVenn (36). Additionally, ADPRylated proteins that were present in at least two of the three cell lines for luminal and basal were overlapped and presented in a Venn diagram generated with bioVenn (36).

#### Examining the overlap of luminal and basal substrates with substrates reported in literature

Venn diagram was created with bioVenn showing the overlap of the PARP1 substrate proteins in our study with previously published work from Zhen *et al.* (2017), Anagho-Mattanovich *et al.* (2025), and Gibson *et al.* (2016) (34,37,38).

#### ADPRylation sites relative to previously published ADPRylation sites

Histograms were generated using R (ver 4.4.1) examining the overlap of our PARP1-mediated ADPRylated sites with previously published works of Gibson et al. (34).

#### Amino acid enrichment around ADPRylation sites

Sequences spanning ± 8 amino acids around all ADPRylated sites for PARP1 were analyzed using a custom *Python* (ver 3.12) script to quantify amino acid frequencies at each position and stacked bar plot was generated to visualize the distribution of amino acids surrounding these ADPRylated sites. Additionally, a sequence logo was generated using the logomaker package highlighting amino acids with frequencies ≥ 10%.

#### ADPRylation sites relative to other post-translational modifications

Known post-translational modifications (phosphorylation, ubiquitylation, acetylation, methylation, and sumoylation) were obtained from the iPTMnet database, which is a curated database sourced from HPRD, PhosphoSitePlus, UniProt, etc. (39). The sites of ADPRylation identified in this study alongside an aspartate/glutamate ratio-normalized random control set were analyzed relative to other PTMs using the iPTMnet database. The PTM with the closest distance to an ADPRylated site or random control were kept for analysis and visualized using histograms in R (ver 4.4.1). Fisher’s exact test was utilized to examine the differences between PTM site counts within ± 5 amino acids of the ADPR site and the random control. Significance was defined with a p-value < 0.05.

#### Gene Ontology analyses

Gene ontology analyses were carried out using the DAVID (Database for Annotation, Visualization, and Integrated Discovery) tool (40). The input was the PARP1 substrate proteins found to be ADPRylated in luminal cell lines, basal cell lines, and the overlap of both luminal and basal. Heatmap was created in R (ver 4.4.1) using the *pheatmap* package showcasing the enrichment scores, log (p-value), of the top GO terms associated with biological processes.

#### Identification of ADPRylated transcription factors

Transcription factors that are ADPRylated across all six breast cancer cell lines and subtype-specific were identified using the JASPAR database (41).

#### Identification of ADPRylated chromatin remodeling complexes, histone-modifying complexes, and linker and core histones

ADPRylated proteins were cross-referenced with published subunit list of complexes. Subtype-specific subunits as well as subunits present across all six breast cancer cell lines were highlighted.

#### Data visualization and statistics

Heatmaps were generated in R (ver 4.4.1) using the *pheatmap* package to showcase the protein abundance patterns of both the total proteome and ADPRylated proteome. Normalized protein abundance values were log_2_ transformed and scaled by cell line to enable comparison across cell lines. Volcano plots were constructed in R (ver 4.4.1) using the *ggplot2* package. Protein-level abundances for ADPRylation and total proteome across the two groups: luminal (MCF-7, T-47D, ZR-75) and basal (HCC70, MDA-MB-231, MDA-MB-468) were derived from the mass spectrometry data. For each protein and replicate, we computed *r = log_2_(ADPR/Total).* To minimize outlier influence, we selected two cell lines in each group with the smallest absolute deviation from the group median of *r*. The top 2 values were used for downstream analysis. For each protein, log_2_ fold change (basal vs luminal) was the difference of the group means from the selected pairs. Significance was tested with a two-sided Welch’s t-test on the selected pairs (n = 2 versus n =2). Proteins were labeled “More ADPRylated in Basal/TN” or “More ADPRylated in Luminal” if fold change > 1.5 and p < 0.05, otherwise they were termed as non-significant. Volcano plot shows log_2_FC(x) vs -log10p(y) with dashed reference lines at specified thresholds.

### RNA sequencing and data analysis

#### Generation of RNA-seq libraries

T-47D and MDA-MB-468 cells were treated with DMSO or Olaparib at 20 μM for 6 hours. Total RNA was isolated using the RNeasy kit (Qiagen, 74106) according to the manufacturer’s instructions. The NEBNext Ultra II Directional RNA Library Prep (NEB, E7765L) in conjunction with Poly(A) mRNA Magnetic Isolation Module kit (NEB, E7490L) was then used to prepare libraries according to the manufacturer’s instructions. The RNA-seq libraries were subjected to QC analyses [final library yields as assessed by Qubit 2.0 (Invitrogen) and size distribution of the final library DNA fragments as assessed by TapeStation (Illumina)] and sequenced using an Illumina HiSeq 2000.

#### Analysis of RNA-seq data

Raw FASTQ files underwent quality control with FastQC (Babraham Bioinformatics, v0.11.x) (1). Reads were aligned to the human reference genome (GRCh38/hg38) using TopHat2 (2) v2.0.12 with Bowtie2 v2.1.0 (3) as the aligner backend (key options: --keep-fasta-order --no-coverage-search --library-type fr-firststrand -G <GENCODE GTF> --transcriptome-index <IDX> -g 10). Alignment files were processed with SAMtools v0.1.19 (4). Transcriptome assembly was performed with Cufflinks v2.2.1 (5), transcripts were merged with Cuffmerge, and differential expression was assessed with Cuffdiff using default parameters.

#### Transcriptome data analyses

Genes were called differentially regulated if they satisfied both a fold-change threshold (FC ≥ 1.5 for upregulated; FC ≤ 0.67 for downregulated) and a significance cutoff of p ≤ 0.05.

#### Gene Ontology analyses

Gene ontology analyses were conducted using the Database for Annotation, Visualization, and Integrated Discovery (DAVID) Bioinformatics Resources tool (42,43). Log_10_ p-values and percent of targets in each GO term were presented for top 10 relevant terms.

### Immunoprecipitation (IP)

T-47D and MDA-MB-468 cells were seeded at ∼5 × 10^6^ cells per 15 cm diameter plate and cultured. The cells were collected and resuspended in IP Lysis Buffer (25 mM Tris-HCl pH 7.5, 450 mM NaCl, 2 mM MgCl2, 1 mM CaCl_2_, 0.2% Triton X-100, with freshly added 40 U/mL micrococcal nuclease, 250 nM ADP-HPD, 10 µM PJ34, and 1x complete protease inhibitor cocktail), incubated for 30 min at 4°C with intermittent vigorous vortexing. The micrococcal nuclease reaction was stopped by adding 5 mM EDTA. After centrifugation at 20,000 x g for 10 min at 4°C, the supernatant was collected, and the concentration was determined using the BCA protein assay. For each cell line, 1 mg of nuclear extract was mixed with two volumes of Dilution Buffer (25 mM Tris-HCl pH 7.5, 2 mM MgCl_2_, with freshly added 250 nM ADP-HPD, 10 µM PJ34, and 1x complete protease inhibitor cocktail), and incubated with TFAP2A antibody conjugated to agarose (Santa Cruz, sc-12726 AC; RRID:AB_667767) overnight at 4°C. The agarose beads were then washed three times in Wash Buffer (25 mM Tris-HCl pH 7.5, 150 mM NaCl, 2 mM MgCl_2_, 0.2 mM EDTA) for 10 min at 4°C with constant mixing. The beads were then heated to 95°C for 5 min in 2x SDS-PAGE loading buffer to release the bound proteins. The immunoprecipitated material was subjected to Western blotting.

### Chromatin immunoprecipitation (ChIP)

#### Chromatin immunoprecipitation

For ChIP assays, cells were cross-linked with 1% formaldehyde in PBS for 10 minutes at 37^°^C, quenched by the addition of 125 mM glycine, and incubated 5 minutes at 4^°^C. The cross-linked cells were collected in ice-cold PBS and pelleted by centrifugation. The cells were then resuspended by gentle mixing by pipetting in ice-cold Farnham Lysis Buffer (5 mM PIPES pH 8.0, 85 mM KCl, 0.5% NP-40) with freshly added 1 mM DTT and 1x protease inhibitor cocktail. The supernatant was removed, and the crude nuclear pellet was collected by centrifugation and resuspended in SDS Lysis Buffer (50 mM Tris-HCl pH 7.9, 1% SDS, 10 mM EDTA) with freshly added 1 mM DTT and 1x protease inhibitor cocktail. After a 10-minute incubation on ice, the lysate was sheared by sonication using a Bioruptor (Diagenode) to generate chromatin fragments of approximately 250 bp in length. The sheared chromatin was clarified by centrifugation and the diluted ten-fold in ChIP Dilution Buffer (20 mM Tris-HCl pH 7.9, 0.5% Triton X-100, 2 mM EDTA, 150 mM NaCl) with freshly added 1 mM DTT and 1x protease inhibitor cocktail. The lysate was pre-cleared with equilibrated protein A-agarose beads and subjected to immunoprecipitation reactions with IgG or an antibody against TFAP2A at 4^°^C overnight.

The immunoprecipitates were collected by incubation with BSA-blocked protein A-agarose beads for 2 hours at 4^°^C with gentle mixing. After incubation, the beads were washed on ice once each with (1) Low Salt Wash Buffer (20 mM Tris-HCl pH 7.9, 2 mM EDTA, 125 mM NaCl, 0.05% SDS, 1% Triton X-100, 1x protease inhibitor cocktail), (2) High Salt Wash Buffer (20 mM Tris-HCl pH 7.9, 2 mM EDTA, 500 mM NaCl, 0.05% SDS, 1% Triton X-100, 1x protease inhibitor cocktail), (3) LiCl Wash Buffer (10 mM Tris-HCl pH 7.9, 1 mM EDTA, 250 mM LiCl, 1% NP-40, 1% sodium deoxycholate, 1x protease inhibitor cocktail), and (4) 1x Tris-EDTA (TE). The beads were then subjected to a final wash with 1x TE at room temperature. They were then collected by centrifugation, resuspended in ChIP Elution Buffer (100 mM NaHCO3, 1% SDS), and incubated on end-over-end rotator for 15 minutes at room temperature to elute the ChIPed DNA. The ChIPed DNA was de-crosslinked by adding 100 mM NaCl with incubation at 65^°^C overnight. The eluted material was cleared of protein and RNA by adding RNase H and proteinase K, and incubating for 2 hours at 55^°^C. The ChIPed DNA was then extracted with phenol:chloroform:isoamyl alcohol (25:24:1), collected by ethanol precipitation, and dissolved in water. ChIPed DNA was used for library preparation for sequencing.

#### ChIP-qPCR

The ChIPed genomic DNA was subjected to qPCR using gene-specific primers. The immunoprecipitation of genomic DNA was normalized to the input. All experiments were performed a minimum of two times with independent biological replicates.

#### Primers for ChIP-qPCR

The following primers were used:

- SET Forward: 5’-CTTCGCCTTCCCTTCTCTCC-3’
- SET Reverse: 5’-TTTTACTGACTTTGGCCGCC-3’
- FLII Forward: 5’-AGGACAGGCATTCAAGACCA-3’
- FLII Reverse: 5’-AGCTGGGATTATAGGCGTGC-3’

### Data availability statement

Data generated in this study has been included in the manuscript and Supplementary Data files. RNA-Seq data generated were deposited in GEO under accession number GSE308423. Mass spectrometry data generated were deposited in MASSIVE under the ID MSV000099017. Software, scripts, additional data, and other information about the analyses can be obtained by contacting the corresponding author (W.L.K.).

## Results

### Differential sensitivity of breast cancer cell lines to PARP inhibitors

To explore the subtype-specific ADPRylated proteome in breast cancer cells, we selected six well-characterized breast cancer cell lines representing luminal A (MCF-7, T-47D, ZR-75) and basal/triple-negative (HCC70, MDA-MB-231, MDA-MB-468) subtypes (**Fig. 1A**). All six of the cell lines have wild-type (unmutated) *BRCA1* and *BRCA2* genes, but some harbor mutations in *TP53* (the gene encoding the tumor suppressor protein p53) or other DNA damage repair (DDR) proteins (**Fig. 1A**). All of the cell lines exhibited detectable basal levels of PARylation by Western blotting, with MDA-MB-468 showing considerably higher (∼5-fold) levels than the other cell lines (**Fig. 1B**). Upon H O -induced genotoxic stress and DNA damage, PARP1 activity was strongly induced in all lines, and this activity was efficiently suppressed by pretreatment with Olaparib or Talazoparib (**Suppl. Fig. S1, A and B**). Any array of publicly available genomic data, including RNA-seq, GRO-seq, and ChIP-seq for histone modifications, is available for many of these cell lines (44), increasing their utility as models for ‘omic’ analyses.

**Figure 1.**
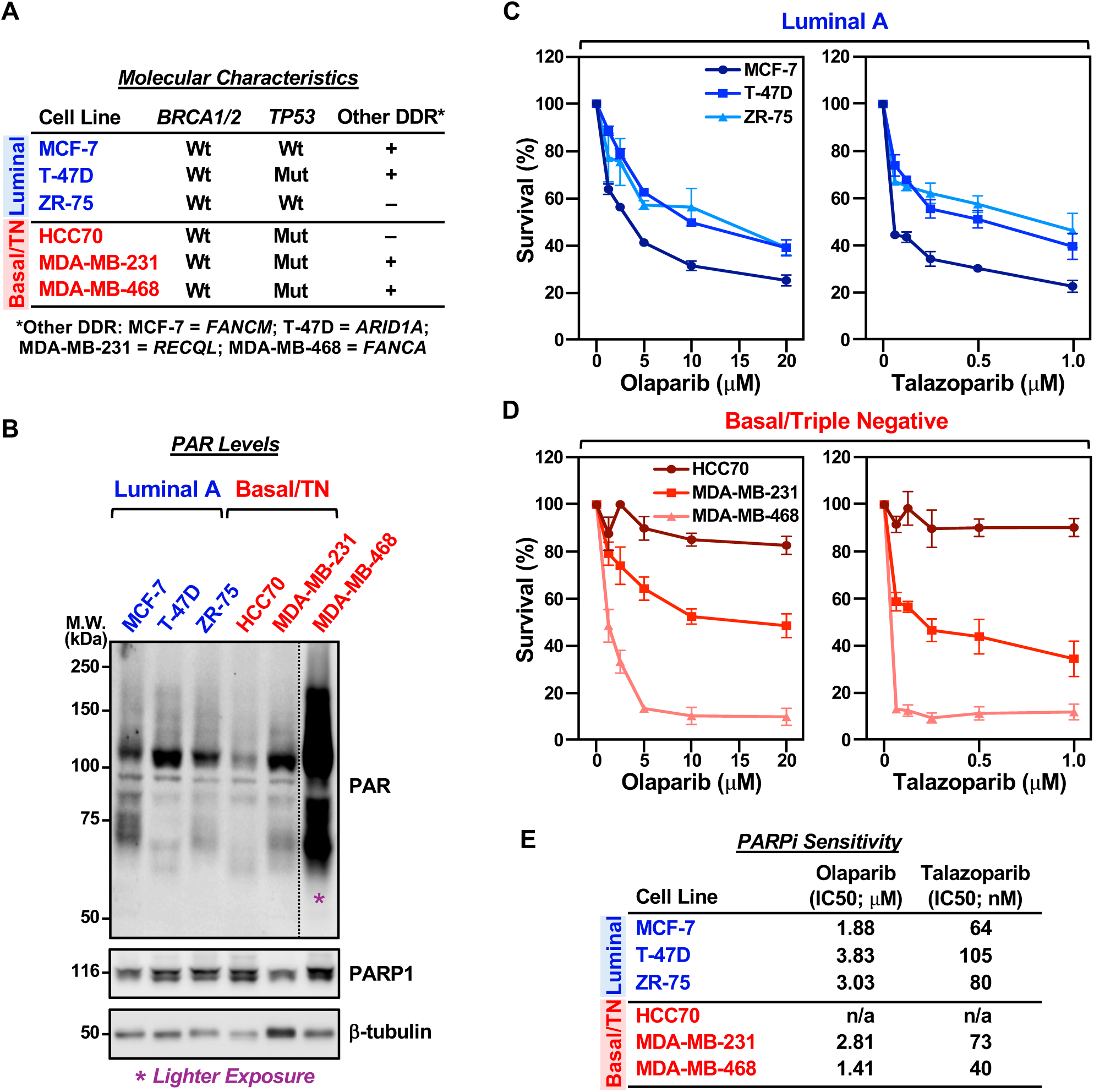
Breast cancer cell lines exhibit different levels of PARP1 sensitivity to PARP inhibitors. **(A)** Table summarizing important molecular characteristics relevant to PARP inhibitor sensitivity of the 6 breast cancer cell lines used in this study, such as mutational status of *BRCA1/2*, *TP53* and other DNA damage repair (DDR) proteins. **(B)** Western blot showing basal levels of ADPRylation (PAR) and PARP1 in whole cell extracts prepared from three luminal A (*blue*) and 3 basal/triple negative (*red*) breast cancer cell lines. β-tubulin was used as a loading control. Molecular weight markers in kilodaltons (kDa) are indicated. **(C and D)** Dose response curves showing increasing concentrations of Olaparib (*left panel*) or Talazoparib (*right panel*) significantly decreasing cell viability in luminal A (C) or basal/triple negative (D) subtype breast cancer cell lines, as assayed by crystal violet cell proliferation assay over a period of 5 days. Each point represents the mean ± SEM; n = 3. **(E)** Table showing the half maximal inhibitory concentration (IC50) values of Olaparib or Talazoparib in luminal A or basal/triple negative breast cancer cell lines, as calculated from survival curves in panels C and D using GraphPad Prism.

In dose–response assays, both Olaparib or Talazoparib decreased cell viability in a concentration-dependent manner for all cell lines, except HCC70, which was insensitive even at the highest dose (20 µM) (**Fig. 1, C and D**). The IC values ranged from ∼1.4 to ∼3.8 µM for Olaparib and 40 to ∼100 nM for Talazoparib, except for HCC70 cells, for which an IC value could not be determined (**Fig. 1E**). Interestingly, MDA-MB-468 cells had the highest levels of basal PARylation and the lowest IC values, and vice versa for HCC70 cells, perhaps suggesting that higher basal ADPRylation is related to greater drug sensitivity as was observed previously in ovarian cancer cells (45). If so, these results suggest that basal PARP1 activity may contribute to differential drug response.

### Characterization of the PARP1-mediated Glu/Asp ADPRylated proteome using a chemical genetics approach coupled with mass spectrometry

To determine the repertoire of PARP1 substrates across the six breast cancer cell lines, we used a chemical biology approach that we developed previously, which allows for the identification of PARP-specific ADPRylation events (33,34). Our NAD^+^ analog-sensitive PARP (asPARP) approach uses an unnatural NAD^+^ analog, 8-Bu(3-yne)T-NAD^+^, which can only bind to PARPs that have a mutation at a specific “gatekeeper” residue in their catalytic domain. The mutations create a “hole” in the PARP catalytic domain that can be filled by an extra chemical moiety (“bump”) on the NAD^+^ analog (**Fig. 2A**), conceptually similar to the approach developed by Shokat and colleagues for kinases (46). The addition of a “clickable” moiety (i.e., alkyne) on the NAD^+^ analog increases utility by providing a chemical handle that can be clicked to azide in solid substrates (e.g., agarose-azide), biotin (biotin-azide) or fluorophores [e.g., tetramethylrhodamine (TAMRA)-azide] (33,34) (**Fig. 2B**). We validated this system for PARP1 in vitro using recombinant wild-type PARP1 (Wt) or asPARP1 (as) (**Suppl. Fig. S2**), with efficient automodification of PARP1 only observed with 8-Bu(3-yne)T-NAD^+^, not with native NAD (**Fig. 2C**).

**Figure 2.**
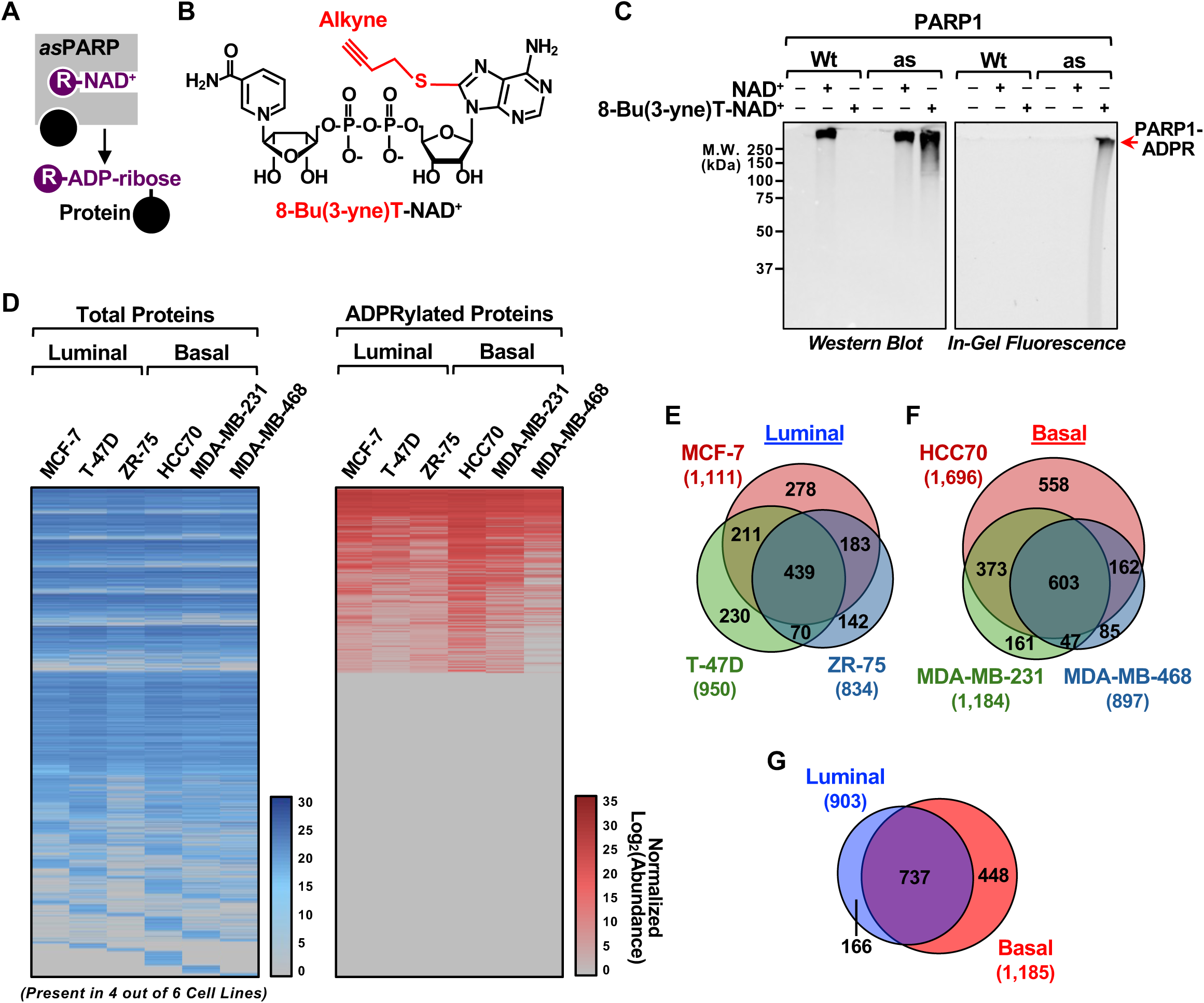
Comprehensive identification of ADPRylated substrates using asPARP1 approach. **(A and B)** Schematic illustrating NAD^+^ analog sensitivity of asPARP1. R indicates the unnatural chemical moiety added to NAD^+^ (A). Chemical structure of the bifunctional NAD^+^ analog 8-Bu(3-yne)T-NAD^+^ with the “clickable” analog sensitivity-inducing, alkyne-containing R group highlighted in red (B). **(C)** In vitro auto-modification assays using purified PARP1 wild-type (Wt) or analog-sensitive (as), and NAD^+^ or 8-Bu(3-yne)T-NAD^+^. Western blot (*left panel*) and in-gel fluorescence assay (*right panel*) conjugated to azido-TAMRA after labeling reactions with 8-Bu(3-yne)T-NAD^+^ in the presence of asPARP1, showing the levels of ADPRylation on PARP1. Ponceau S staining was used to confirm equal loading of material. Molecular weight markers in kilodaltons (kDa) are indicated. **(D)** Heatmaps showing abundance of total proteome (*left panel;* n = 4926) and ADPRylated proteome (*right panel*) in luminal and basal breast cancer cell lines identified by mass-spectrometry. Normalized protein abundance values were log_2_ transformed and scaled by cell line to enable comparison across cell lines. **(E-G)** Venn diagram showing the overlap of ADPRylated substrates mediated by asPARP1 in 3 luminal (E) or 3 basal/triple-negative (F) subtype breast cancer cell lines as assessed by mass-spectrometry. The ADPRylated substrates shared in at least two of the three cell lines for luminal and basal were overlapped and presented in a Venn diagram (G).

We then applied the asPARP1 platform to nuclear extracts from each of the six breast cancer cell lines, followed by: (1) clicking to agarose-azide, (2) stringent washing, (3) elution of peptides not covalently linked to the agarose via an ADPR linkage (for peptide identifications; IDs), (4) another round of stringent washing, and (5) elution using hydroxylamine (NH_2_OH), which cleaves the ester linkage between ADPR and acidic (i.e., Glu, Asp) residues (35). We identified ADPRylated substrates by hydroxylamine cleavage (for ADPR site IDs). We then performed liquid chromatography-tandem mass spectrometry (LC-MS/MS) of the trypsin-eluted peptides, which yielded the total ADPRylated proteome for any ADPRylated residue, and the hydroxylamine-eluted peptides, which yielded the sites of Glu and Asp ADPRylation (**Supplemental Data D1**). We also performed LC-MS/MS of the trypsin-digested input nuclear extracts to define the total proteome (**Supplemental Data D2**). A comparison of the total and ADPRylated proteomes revealed that about 36% of the detectable proteome across the six cell lines was ADPRylated (**Fig. 2D**).

This strategy yielded a robust catalog of thousands of ADPRylated peptides with good replicate reproducibility (∼50-80%) and hundreds of ADPR sites identified across the six cell lines (**Table 1; Supplemental Data D1**). Filtering criteria [i.e., Log_2_[Experimental/Control] > 1; FDR < 1%) were applied to define a stringent substrate set (**Table 1**). Hundreds of PARP1 substrates were shared within subtypes, with 439 proteins common to all three luminal lines and 903 proteins observed in at least two luminal cell lines (**Fig. 2E**) and 603 proteins common to all three basal lines and 1,185 proteins observed in at least two basal cell lines (**Fig. 2F**). Approximately 38% (448/1,185) of the high-confidence ADPRylated proteins were unique to the basal cell lines and ∼18% were unique to the luminal cell lines (**Fig. 2G**). We also observed considerable overlap of the ADPRylated proteins and ADPR sites that we identified with proteins and sites from other published data sets from breast cancer cells (37,38) and cervical cancer cells (34) (**Suppl. Fig. S3, A and B**). Additionally, we observed overlap with some PARP1, PARP2, and PARP3 ADPR sites in a site-to-site comparison (**Suppl. Fig. S3, C through E**). Overall, this catalog of subtype-specific and common PARP1-mediated ADPRylation events defines a broad substrate landscape across breast cancer cells, providing a resource for downstream analyses.

**Table 1.**
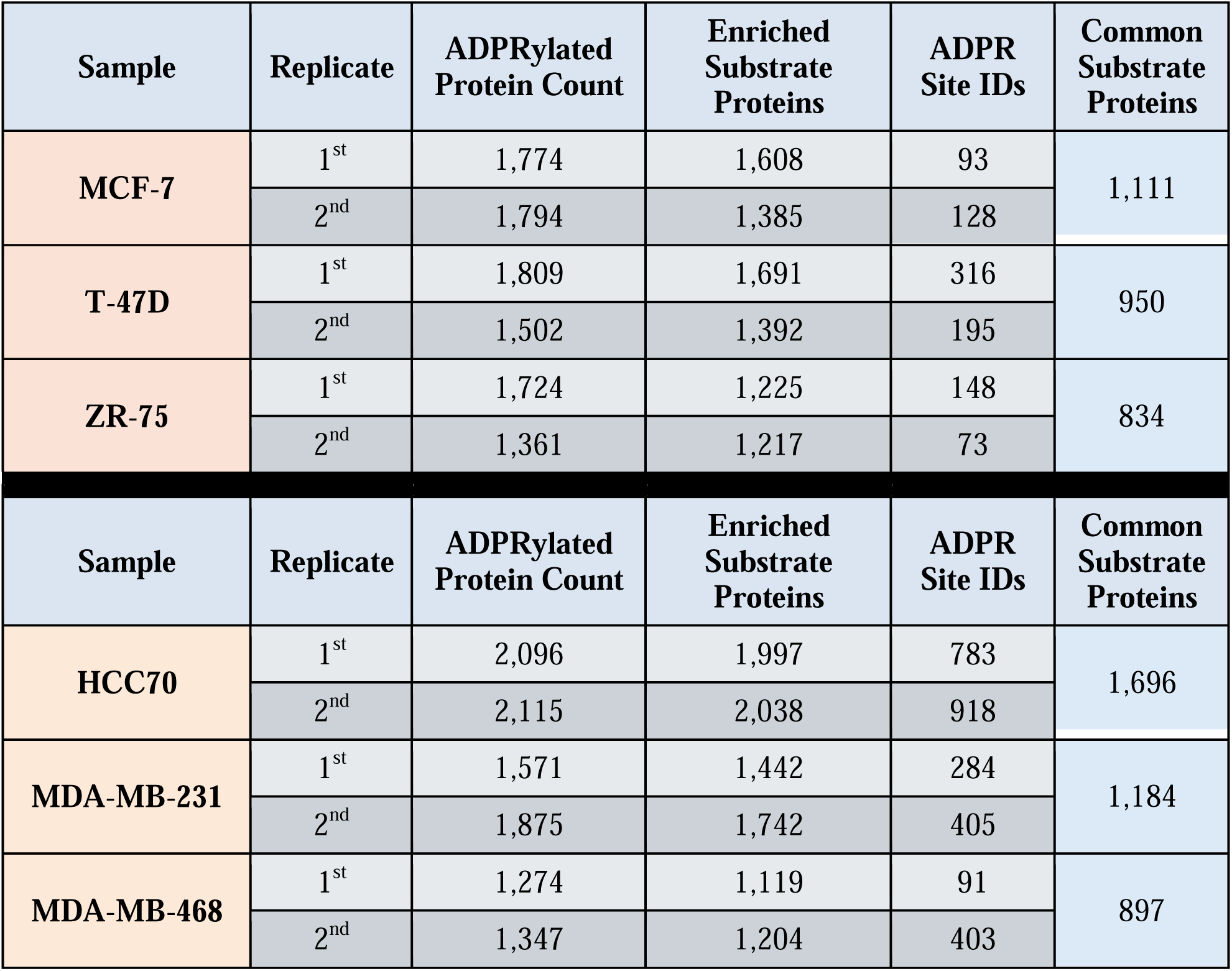
Determination of the PARP1 ADPRylated proteome in breast cancer cell lines. asPARP1 mass spectrometry data from MCF-7, T-47D, ZR-75, HCC70, MDA-MB-231, and MDA-MB-468 cell lines. Nuclear extracts from each sample were subjected to the asPARP1 approach. Enriched Substrates were determined as: Log_2_(Experimental/Control) >1. Experimental = ADPRylated proteins. Control = reaction without asPARP1.

### Sequence-level analyses reveal distinct features of ADPRylation sites

We performed motif analyses of the PARP1-mediated Glu/Asp ADPR sites, which revealed an enrichment of glutamate residues at the site of modification, favored by ∼10 to 1 over aspartate residues (**Fig. 3A**). The enrichment of glutamate and proline residues adjacent to or near the site of ADPRylation is reminiscent of features described previously for PARP1-mediated ADPR sites (34) (**Fig. 3A**). In additional analyses, we mined published databases of known PTMs (e.g., phosphorylation, ubiquitylation, acetylation, methylation, sumoylation), aligned them, and centered them on the sites of ADPRylation that we identified herein. As we described previously (34), we observed a significant enrichment of phosphorylation sites adjacent to or near the site of ADPRylation (**Fig. 3, B and C**), which fits well with the functional interplay between ADPRylation and phosphorylation that we have characterized previously (34,47,48). We also observed a significant enrichment of ubiquitylation sites adjacent to or near the site of ADPRylation (**Fig. 3, D and E**), with ubiquitylation occurring at sites of ADPRylation evident (**Fig. 3D**, see the arrow). The latter observation is consistent with the recent identification of a hybrid ADPR-ubiquitin modification (49–53). Finally, we observed a significant enrichment of acetylation sites, but not methylation or sumoylation sites, adjacent to or near the site of ADPRylation (**Suppl. Fig. S4, A-C**). Collectively, these results show that our asPARP approach robustly and faithfully identifies specific sites of ADPRylation mediated by a specific PARP family member, PARP1.

**Figure 3.**
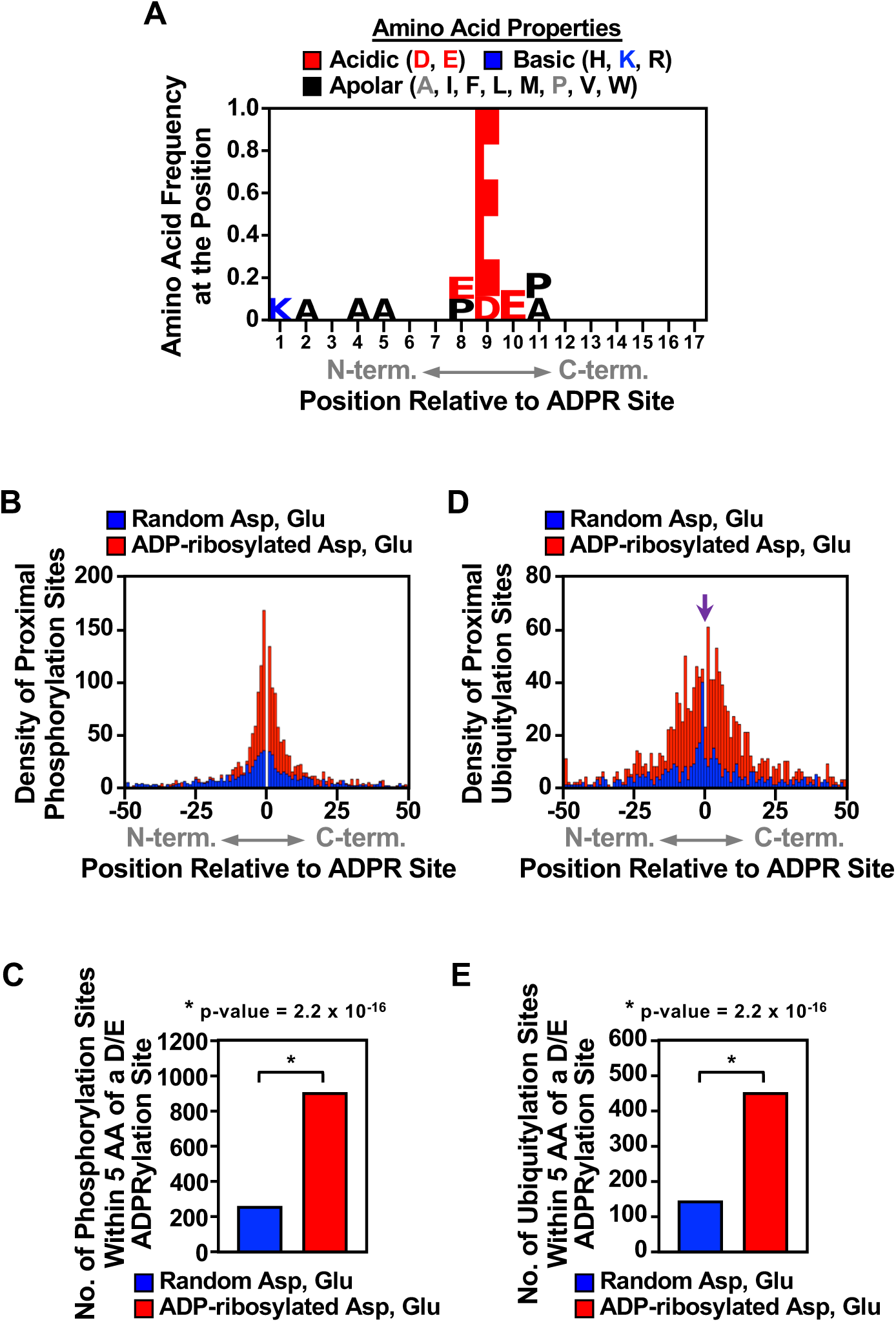
Sequence-level analyses reveal distinct features of ADPRylation sites. **(A)** Enriched amino acid sequences ± 8 residues on either side of the identified PARP1 ADPRylation sites. Different colors indicate different amino acid properties (acidic, basic, or apolar). **(B and D)** Histogram of the 2D relationship between the nearest incidence of known phosphorylation (B) or ubiquitylation (D) sites within ± 50 amino acids of the ADPRylation sites (Asp and Glu) identified herein. A random set of Asp and Glu residues were used as control. In (C), the arrow indicates overlap between the ADPRylation sites and ubiquitylation sites. **(C and E)** Bar graphs showing quantification of phosphorylation (C) or ubiquitylation (E) sites within 5 amino acids of the ADPRylated site. A random set of Asp and Glu residues were used as control. Fisher’s exact test was utilized to examine differences and p-values are noted.

### Gene ontology analyses suggest that distinct biological processes are regulated by PARP1-mediated ADPRylation

We next interrogated the functional significance of the identified substrates using Gene Ontology (GO) analysis. Luminal-, basal-, and common PARP1 substrates were analyzed separately, revealing enrichment in distinct functional categories (**Fig. 4A**). Common substrates were enriched in all of the major ontological categories identified (i.e., RNA splicing, chromatin, transcription, and translation), while luminal-specific substrates were enriched in proteins involved in RNA splicing, chromatin, transcription, and translation and basal-specific substrates were enriched in proteins involved in RNA splicing, chromatin, and translation (**Fig. 4A**). Interestingly, we observed only a limited enrichment of proteins involved in DNA damage responses and DNA repair, limited to common and basal-specific substrates (**Fig. 4A**). The enrichment across all cell types of proteins involved in chromatin and transcription is reflected in the enrichment of ADPRylated proteins in chromatin remodeling complexes (CRCs), histone-modifying enzymes (HMEs), linker and core histones, and transcription factors (TFs) (**Fig. 4, B and C; Suppl. Data D3**). Together, these results illustrate the broader functions of PARP1 based on its substrates and suggest that PARP1 engages core nuclear processes across subtypes, while controlling subtype-specific transcriptional and chromatin regulatory pathways.

**Figure 4.**
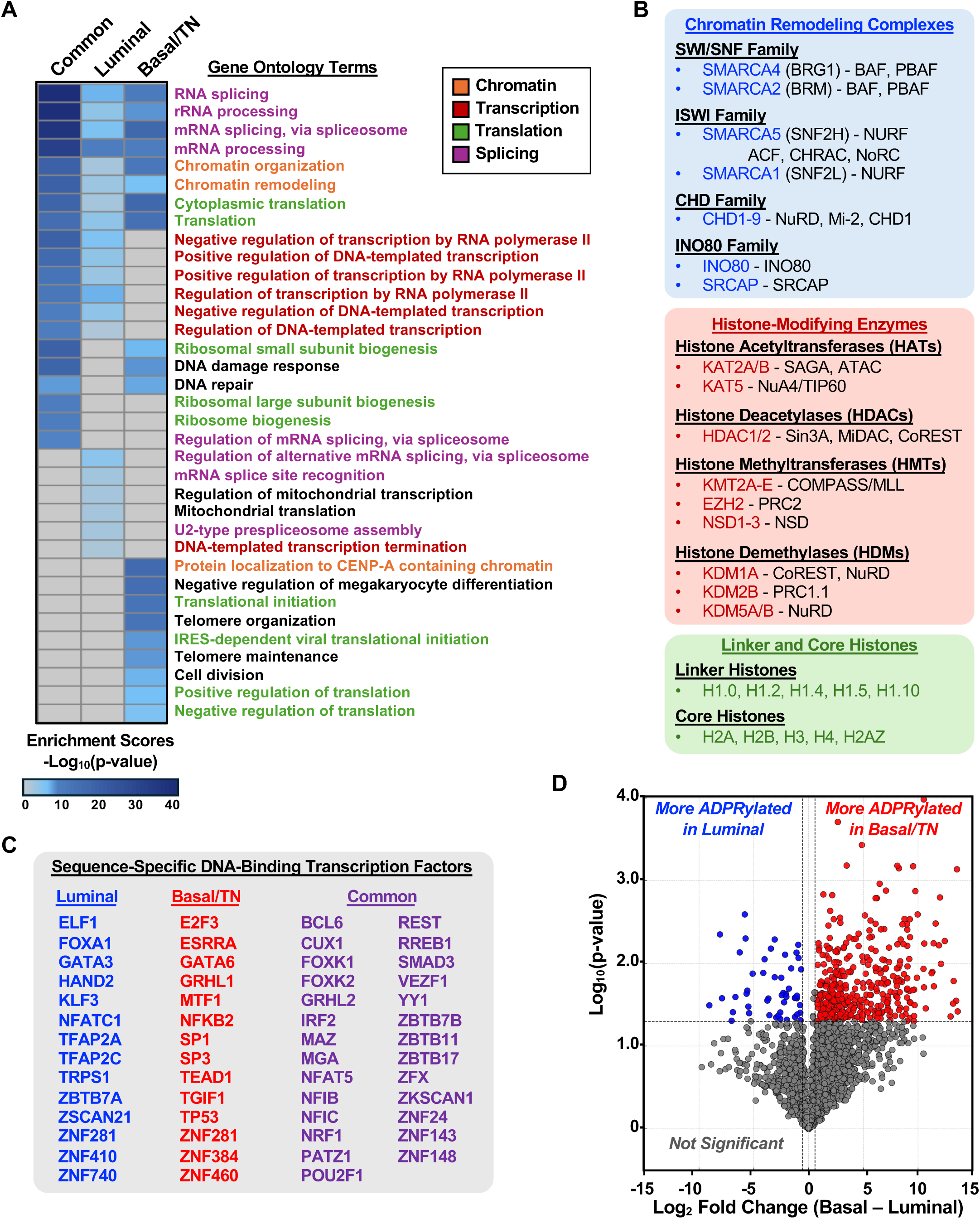
Gene ontology reveals distinct and common biological processes modulated by PARP1 activity in luminal versus basal subtype breast cancer cell lines. **(A)** Heat map of gene ontology (GO) analysis using the DAVID tool from PARP1 substrates identified in Figure 2G. Common (737 proteins), luminal-specific (166 proteins), and basal-specific (448 proteins) substrates were each analyzed for relevant GO terms. Heat map indicates significance of each term based on p-value. Grey color indicates term is not present in the category. Terms are color coded to indicate broader categories of terms related to chromatin, transcription, translation and splicing. **(B)** Categories of identified PARP1 substrates that are part of chromatin remodeling complexes and histone-modifying enzymes, or linker and core histones. The PARP1 substrates are shown in blue, red, or green, and the complex they belong to are listed in black. **(C)** List of sequence-specific DNA-binding transcription factors identified as luminal- or basal-specific substrates of PARP1. Common substrates for both luminal and basal subtypes are also listed. Transcription factors were identified using the JASPAR database. **(D** Volcano plot summarizing differences in ADPR between luminal and basal/triple negative cells normalized to total proteome. For each protein, log (ADPR/Total) was computed per replicate, and the two most concordant replicates per group were used for Welch’s t-test (two-sided). Points are colored blue (more ADPRylated in luminal) or red (more ADPRylated in basal/triple negative) when p < 0.05 and |log FC| > log (1.5); non-significant points are grey. Vertical and horizontal dashed lines mark fold-change and p-value cutoffs, respectively.

Based on the strong connection between nuclear ADPRylation by PARP1 and transcriptional regulation, we performed additional analyses on sequence-specific DNA binding TFs (**Fig. 4C; Suppl. Data D3**). This choice was also informed by our previous studies demonstrating the regulation of TFs, including C/EBPβ and STAT1α, by PARP1-mediated site-specific ADPRylation (47,54). To explore the subtype-specific ADPRylation of TFs more quantitatively, we devised a metric to assess the level of TF ADPRylation standardized to the level of TF protein, which is reflected in the volcano plot in **Fig. 4D**, with specific TFs shown in **Fig. 4C**. These results indicate that TFs are major substrates of PARP1 in breast cancer cells and that the ADPRylation of many of these TFs occurs in a subtype-specific manner, providing a potential regulatory mechanism for subtype-specific gene regulation.

### PARP1 inhibition differentially modulates gene expression programs in luminal versus basal breast cancer cells

Based on the subtype-specific ADPRylation of TFs by PARP1 in luminal versus basal/triple negative cell lines, we hypothesized that these cells would also exhibit subtype-specific gene expression outcomes in response to PARP inhibitor treatment. To test this, we performed RNA-seq in representative luminal (T-47D) and basal (MDA-MB-468) breast cancer cell lines treated with Olaparib (20 μM, 6 hr). This treatment resulted in broad transcriptional changes, with both up- and downregulated genes in each subtype (**Fig. 5, A and B**). GO analysis revealed distinct biological programs in response to Olaparib treatment in the two cell lines. T-47D cells were enriched in terms related to cell division and cell adhesion (**Fig. 5C**). In contrast, MDA-MB-468 cells were enriched in terms related to transcription and proliferation, although both cell lines showed enrichment of terms related to core transcriptional functions (**Fig. 5D**), but with subtype-specific responses (**Fig. 5, D and E**). Thus, inhibition of PARP1 catalytic activity remodels gene expression in both luminal and basal breast cancer cells, but the underlying biological processes are subtype-specific. In fact, we observed little overlap (<10%) in the Olaparib-affected genes between the two cell lines (**Fig. 5, A and E**)

**Figure 5.**
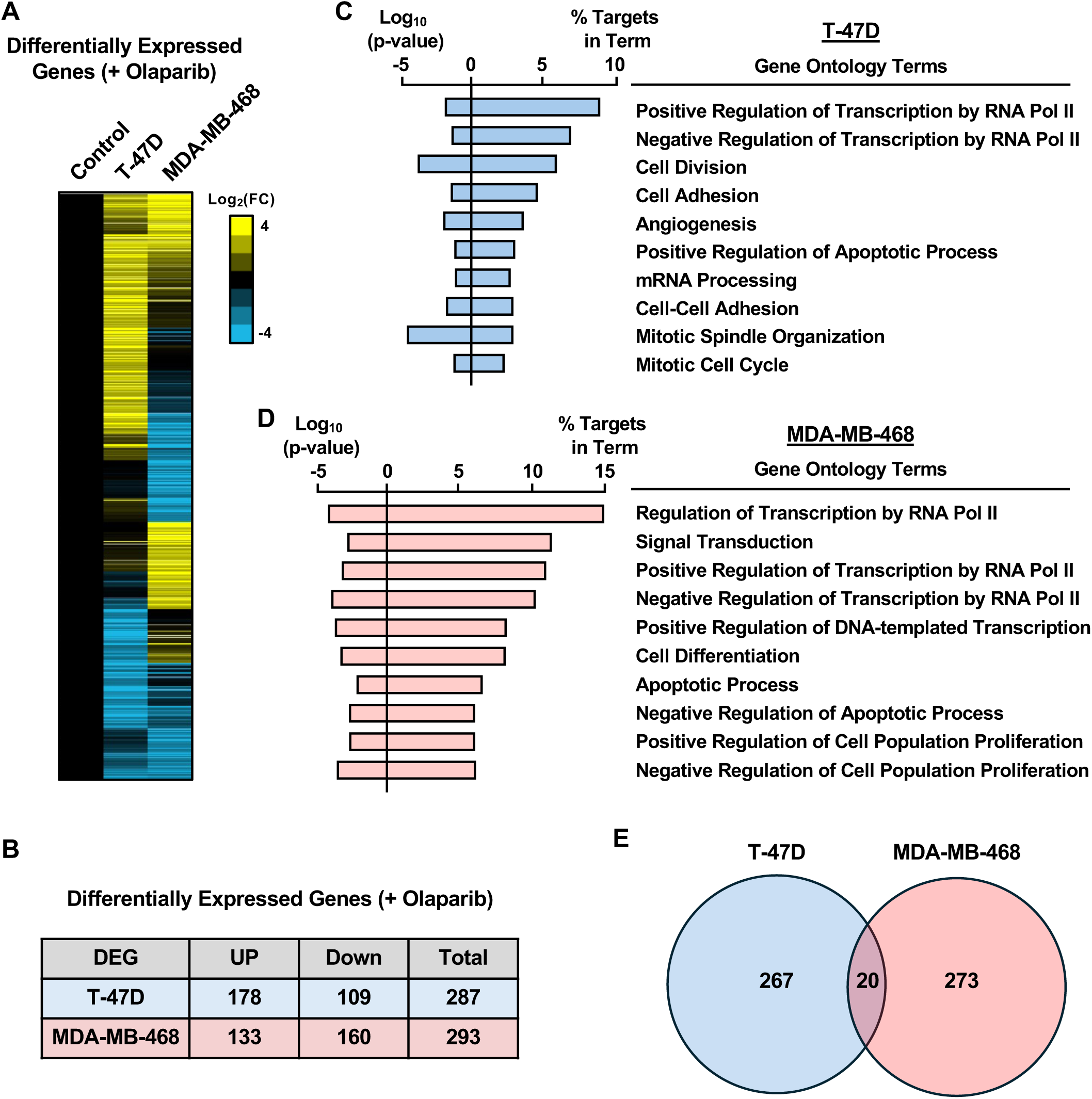
Gene expression reveals distinct biological processes modulated by PARP1 inhibition in luminal versus basal subtype breast cancer cell lines. **(A and B)** Heat map of RNA-seq data showing the differentially expressed genes in response to Olaparib (20 μM for 6 hours) treatment in T-47D and MDA-MB-468 cell lines (A). Yellow indicates upregulated and blue indicates downregulated genes (log_2_ fold-change). Table summarizing the number of up- and downregulated genes (B). **(C and D)** Gene ontology terms from DEGs in response to Olaparib treatment in T-47D (C) and MDA-MB-468 (D). Log_10_ p-values and percent of targets in each GO term were presented for top 10 relevant terms. **(E)** Venn diagram showing the overlap of DEGs in response to Olaparib in T-47D and MDA-MB-468.

### PARP1-mediated ADPRylation regulates TFAP2A function in a breast cancer subtype-specific manner

Based on our results pointing to the subtype-specific regulation of TF ADPRylation by PARP1 as a potential driver of breast cancer cell biology, we sought to confirm key aspects of this hypothesis with a specific example. We mined our proteomics data for a TF with the following properties: (1) broadly expressed across breast cancer cell lines, (2) known cancer-related functions, (3) links to mammary biology and breast cancer, and (4) evidence for subtype-specific ADPRylation from our proteomics data. We chose to focus on TFAP2A, a member of the AP-2 transcription factor family (AP-2 family), which is characterized by an amino-terminal transactivation domain, as well as a carboxyl-terminal helix-span-helix domain, which together with a central basic region, mediates dimerization and DNA binding (55,56). TFAP2A has been implicated in the biology of cancers, including breast cancers (56–60). Moreover, a previous study has shown that it can be PARylated by PARP1 (61). Together, these features make TFAP2A an excellent candidate for proof-of-concept experiments leading from our proteomic data sets to functional outcomes.

We observed variable levels of TFAP2A expression in five out of the six breast cancer cell lines that we tested, with the exception being MDA-MB-231 cells (**Fig. 6A**). We used immunoprecipitation of TFAP2A followed by Western blotting for PAR (IP-Western) in representative luminal (T-47D) and basal (MDA-MB-468) breast cancer cell lines to determine if TFAP2A is PARylated. We observed that TFAP2A was PARylated in T-47D cells, but not in MDA-MB-468 cells (**Fig. 6B**). To assess potential functional consequences, we performed ChIP-qPCR assays at two TFAP2A target gene promoters, *SET* and *FLII*, identified from published ChIP-seq data (62–64) with or without Olaparib treatment. In T-47D cells, PARP1 inhibition reduced TFAP2A binding at the *SET* promoter, whereas it enhanced TFAP2A binding at the *FLII* promoter. In MDA-MB-468 cells, the effects were opposite (enhanced binding of TFAP2A at the *SET* promoter) or attenuated (no effect on the binding of TFAP2A at the *FLII* promoter) (**Fig. 6C**). Together, these results demonstrate that PARP1-mediated ADPRylation modulates TFAP2A promoter occupancy in a subtype-specific manner.

**Figure 6.**
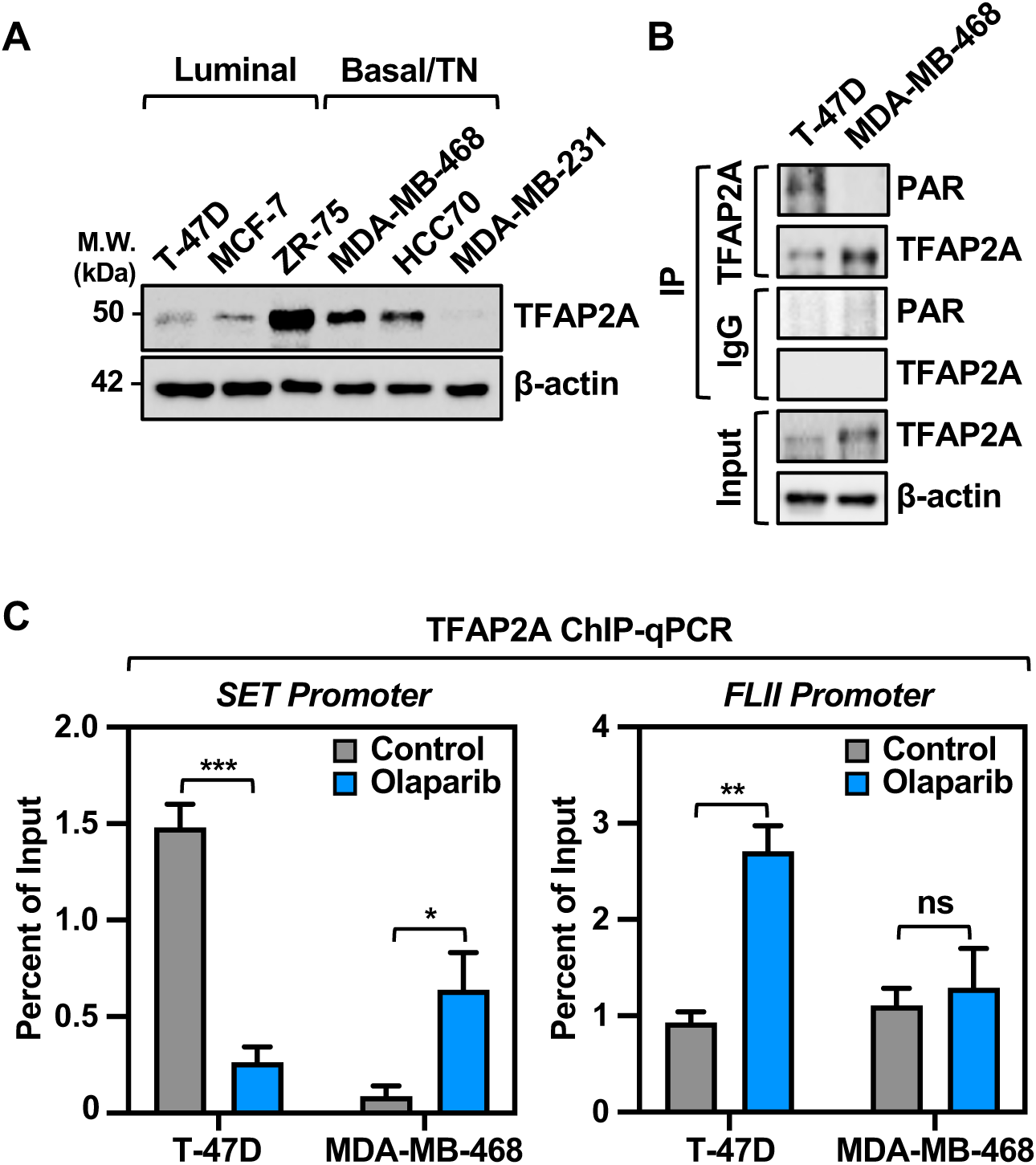
TFAP2A binding to DNA is modulated by ADPRylation status in luminal versus basal subtype breast cancer cell lines. **(A)** Western blot showing expression level of the transcription factor TFAP2A across all six breast cancer cell lines. β-actin was used as a loading control. Molecular weight markers in kilodaltons (kDa) are indicated. **(B)** Western blot showing the T-47D specific ADPRylation of TFAP2A upon immunoprecipitation and blotting for PAR. IgG was used as a control for IP. β-actin was used as a loading control in input. **(C)** Bar graphs from ChIP-qPCR assays showing the enrichment of TFAP2A binding at the *SET* and *FLII* promoters. Differential enrichment upon Olaparib (10 μM for 2 hours) treatment is observed in T-47D (luminal) compared to MDA-MB-468 (basal) cell lines. Each bar represents the mean + SEM; *SET* n = 4, *FLII* n = 3. Bars marked with asterisks are significantly different; unpaired Student’s t-test, * p = 0.05, ** p = 0.01, *** p = 0.001.

## Discussion

Nuclear ADPRylation events, which are mediated primarily by PARP1, have the potential to control a variety of molecular and cellular outcomes. Although the initial studies of ADPRylation were focused on the biochemistry and molecular biology of PARP1 in DNA repair (13), the mechanistic and functional understanding of PARP1 and ADPRylation has grown considerably, including diverse roles in chromatin regulation, transcription, and RNA biology (9,29,30,65). While gene expression profiling has been used to characterize the molecular heterogeneity of cancers, including multiple molecular subtypes of breast cancer (1,2), recent studies have highlighted a growing interest in using ADPRylation profiling in breast cancers in a similar vein (35,37,38,66). In this regard, we have generated new mass spectrometry data sets exploring the ADPRylated proteomes across six breast cancer cell lines representing two prevalent subtypes of breast cancer, luminal and basal/triple negative. Our data sets, which include the ADPRylated proteome, specific sites of Glu/Asp ADPRylation, and corresponding total proteomes, provide a rich resource for the community to interrogate PARP1 signaling in the context of breast cancer heterogeneity.

### New data sets for exploring subtype-specific differences in the PARP1-mediated ADPRylated proteome and therapeutic responses

Our chemical genetics-mass spectrometry platform yielded thousands of ADPRylated proteins and hundreds of sites across six commonly used breast cancer cell lines, providing an extensive catalog of PARP1 substrates that complement previously published data sets (35,37,38,66). Unlike more general approaches for detecting total ADPRylation, our asPARP approach allows assignment of ADPRylation events specifically to PARP1 (33,34). We observed substantial overlap of PARP1 substrates between the luminal and basal subtypes, reflecting shared engagement of core nuclear processes, but we also identified hundreds of subtype-specific substrates. Gene ontology analyses of the PARP1 substrates revealed that while chromatin-, transcription-, and RNA processing-related pathways were enriched across both subtypes, the basal cell lines showed greater enrichment for translation-related pathways, whereas the luminal cell lines displayed stronger connections to chromatin- and transcription-related pathways. These differences suggest that PARP1 activity may reinforce lineage-specific transcriptional programs while maintaining common nuclear functions across subtypes. Notably, DNA damage response and repair pathways were only modestly enriched across the cell lines surveyed under basal growth conditions in our experiments. The utility of these proteomic data sets is amplified by the availability of extensive complementary genomics data for many of these lines (44), enabling integrative analyses of PARP1 function across multiple regulatory layers.

Our results show similarities to other data sets exploring the nuclear ADPRylated proteome using overlapping, but distinct, collections of breast cancer cell lines (35,37,38,66). These studies share the common observation that nuclear ADPRylation is both widespread and context dependent in breast cancer (35,37,38,66). Zhang *et al.* (2013) established the breadth of Glu/Asp ADPRylation across the nuclear proteome (35), while Zhen *et al.* (2017) demonstrated that the proteins modified vary markedly between breast cancer cell lines despite similar PARP1 expression, suggesting subtype-specific signaling heterogeneity (37). Two papers from the Nielsen lab extended these observations to *BRCA1/2* status and acquired PARP inhibitor resistance (38,66). Interestingly, a previous study has shown that the basal activity of PARP1 varies dramatically across a large panel of breast cancer cell lines (67), which we also observed to some extent in our smaller selection of cell lines. Collectively, these results indicate that while ADPRylation is a core nuclear regulatory process, its outputs are highly cell type- and context-specific, with modest changes sufficient to alter therapeutic responses.

### Transcription factors as key substrates of PARP1

The enrichment of transcription factors among PARP1 substrates highlights a central role for ADPRylation in regulating transcriptional networks. We highlighted TFAP2A as an example of a context-dependent substrate, with PARP1-dependent modification altering its promoter occupancy differently in luminal versus basal cells. These findings build on previous work from our lab and others showing site-specific regulation of transcription factors in various biological contexts, such as C/EBPβ (54), STAT1α (47), NFAT (68), and p53 (69). Site-specific ADPRylation of proteins can (1) alter the biochemical or biophysical properties of the modified protein or (2) create new binding sites for ADPR binding domains to drive protein-protein interactions (9,27). Interestingly, ADPRylation is highly associated with other PTMs, such as phosphorylation, across the proteome [results herein; (34,70,71)], which may also modulate the effects of PARP1-mediated ADPRylation on TFs [e.g., as observed with STAT1α (47)].

Previous mapping of the Glu/Asp ADPRylated proteome across breast cancer cell lines identified a large cohort of proteins exhibiting breast cancer subtype-specific patterns of ADPRylation, including a number of TFs (37). For example, the transcription factor GATA3, which is required for the maintenance of luminal cell differentiation (72), is selectively ADPRylated in ERα+ luminal breast cancer cells (37). Another transcription factor, FOXA1, which is a key mediator of ERα-dependent signaling (73,74), is also selectively ADPRylated in ER+ luminal breast cancer cells (37). The subtype-specific ADPRylation patterns of TFs suggests distinct regulatory mechanisms for ADPRylation in different breast cancer subtypes, as well as a general paradigm by which PARP1 fine-tunes transcription factor activity to shape subtype-specific gene expression.

## Acknowledgements

We thank members of the Kraus lab for their helpful comments and support; We also acknowledge and thank the UT Southwestern Proteomics Core Facility for mass spectrometry (Dr. Andrew Lemoff), the Next Generation Sequencing Core for deep sequencing services (Dr. Ralf Kittler and Vanessa Schmid). This work was supported by grants from the NIH/National Cancer Institute (R01 CA251943 to W.L.K.), NIH/National Institute of Diabetes and Digestive and Kidney Diseases (R01 DK069710 to W.L.K.), UTSW American Cancer Society – Institutional Research Grant (IRG-21-142-16 to C.V.C.), UTSW Cancer Center Support Grant (P30CA142543 to C.V.C.) and funds from the Cecil H. and Ida Green Center for Reproductive Biology Sciences Endowment to W.L.K.

## Authors’ Contributions

S.K.- formal analysis, investigation, validation, visualization, data curation. M.K.- formal analysis, investigation, validation, visualization. P.T.- formal analysis, investigation, validation. Y.D. - formal analysis, investigation, validation. T.N.- data curation, formal analysis. D.H.- formal analysis, investigation, validation, visualization. C.V.C.- formal analysis, validation, visualization, writing- original draft preparation, writing- review and editing. W.L.K.- conceptualization, funding acquisition, project administration, supervision, visualization, writing- original draft preparation, writing- review and editing.

## Note

Supplemental Data can be found with this article online including Supplemental Figures S1 through S4 and Supplemental Data D1 through D3. *[See the Supplemental Figure and Supplemental Data files.]*

