## Supplemental Materials for "Mapping the Subtype-Specific PARP1 ADP-ribosylated Proteome in Breast Cancer Cells"

**Koul *et al.* (2025)**

SUPPLEMENTAL DATA Page

### SUPPLEMENTAL DATA
*[see the Excel files provided]*

**Supplemental Data D1. Characterization of the ADPRylated proteome of six breast cancer cell lines determined by an asPARP1 approach with mass spectrometry.**
The trypsin digested (Protein ID) and hydroxylamine eluted (Site ID) asPARP1 samples were subjected to LC-MS/MS analysis. The "Column Heading" key provides information describing the metrics of each of the LC-MS/MS-identified peptides. The "Tab" key provides details about all of the other worksheets within this spreadsheet.

**Supplemental Data D2. Characterization of the total proteome of six breast cancer cell lines using mass spectrometry.**
Nuclear extracts obtained from each individual cell line were subjected to LC-MS/MS analysis. The ""Column Heading"" key provides information describing the metrics of each of the LC-MS/MS-identified peptides. The ""Tab"" key provides details about all of the other worksheets within this spreadsheet.

**Supplemental Data D3. Transcription factors, histones, histone modifying enzymes, and ATP-dependent chromatin remodeling complexes that are ADPRylated by PARP1 in breast cancer cells.**
ADPRylation was determined using the asPARP1 approach in six different breast cancer cells lines. Each tab highlights cell type-specific and breast cancer subtype-specific, as well as common, substrates. Substrates with site IDs are highlighted in red. Additional information is provided for the histone modifying enzyme complexes and chromatin remodeling complexes that contain specific PARP1 substrates.

### Supplemental Figures

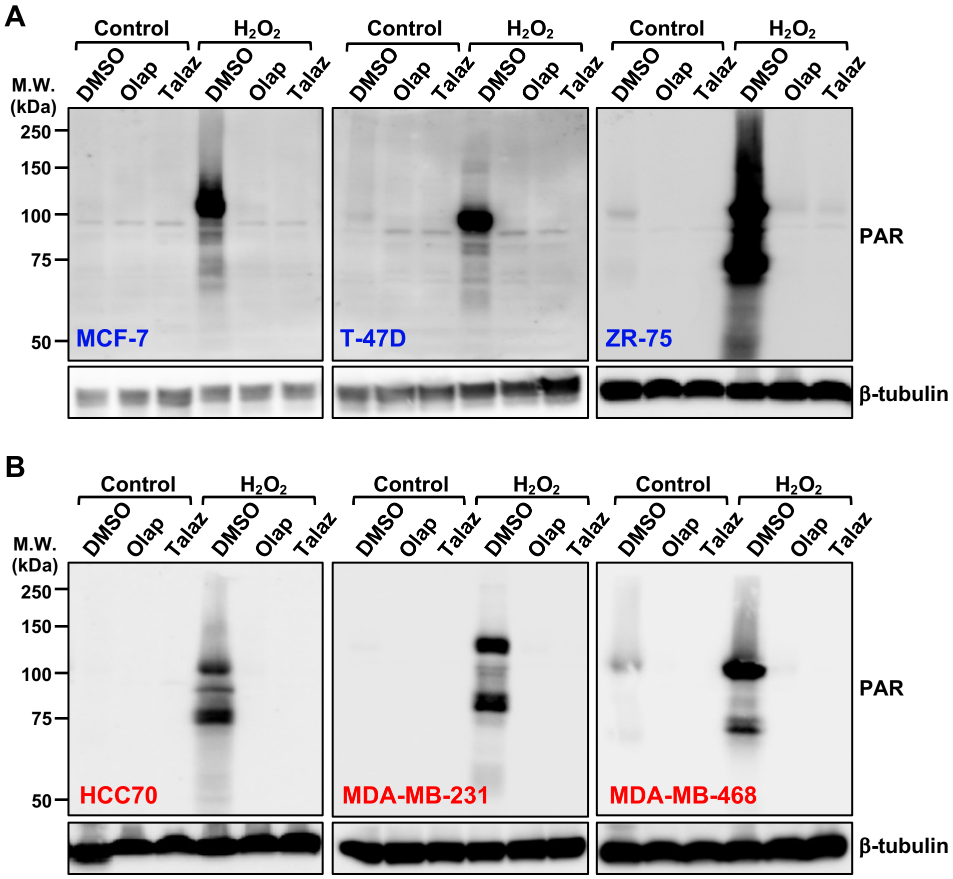

**Figure S1. Breast cancer cell lines exhibit different levels of PARP1 activity.**
**(A and B)** Western blots showing levels of ADPRylation (PAR) with and without treatment with PARP1 inhibitors Olaparib (20 µM) or Talazoparib (1 µM) for 2 hours in luminal (A) or basal/triple negative (B) breast cancer cell lines. The cells were additionally treated with or without $H_2O_2$ (1mM) for 5 minutes and harvested. β-tubulin was used as a loading control. Molecular weight markers in kilodaltons (kDa) are indicated.
*[Related to Figure 1B]*

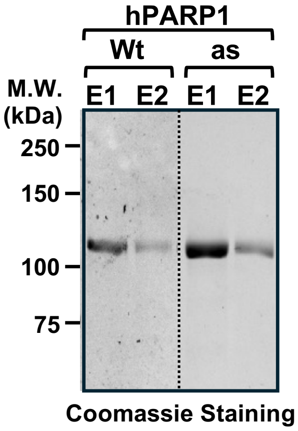

**Figure S2. Purification of asPARP proteins.**
Coomassie blue staining of FLAG-tagged wild-type (Wt) or analog sensitive (as) human recombinant PARP1. Both proteins were expressed in Sf9 using baculovirus and affinity-purified. Molecular weight markers in kilodaltons (kDa) are indicated.
*[Related to Figure 2C]*

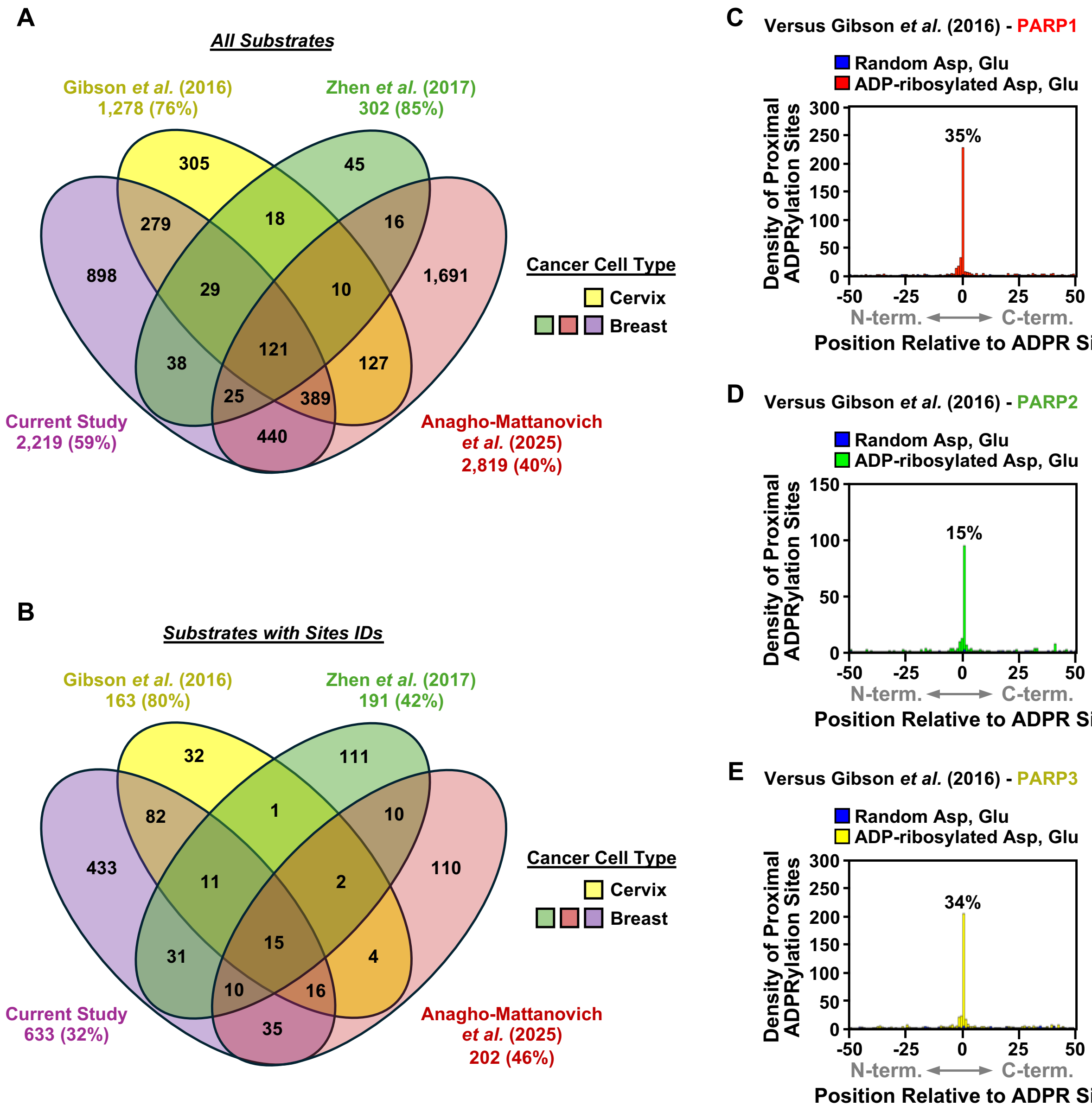

**Figure S3. Comparison of ADPRylation substrates and site IDs across studies.**
**(A and B)** Venn diagrams showing the overlap of total identified ADPRylated substrates (A) and site IDs (B) across three published studies using breast cancer cells [Zhen *et al.* (37) and Anagho-Mattanovich *et al.* (38)] and cervical cancer cells [Gibson *et al.* (34)] and the new data generated herein.
**(C-D)** Histogram of the 2D relationship between previously identified ADPRylation sites from PARP1 (C), PARP2 (D), and PARP3 (E) [Gibson *et al.* (34)] with those sites identified herein.
*[Related to Figure 2E through 2F]*

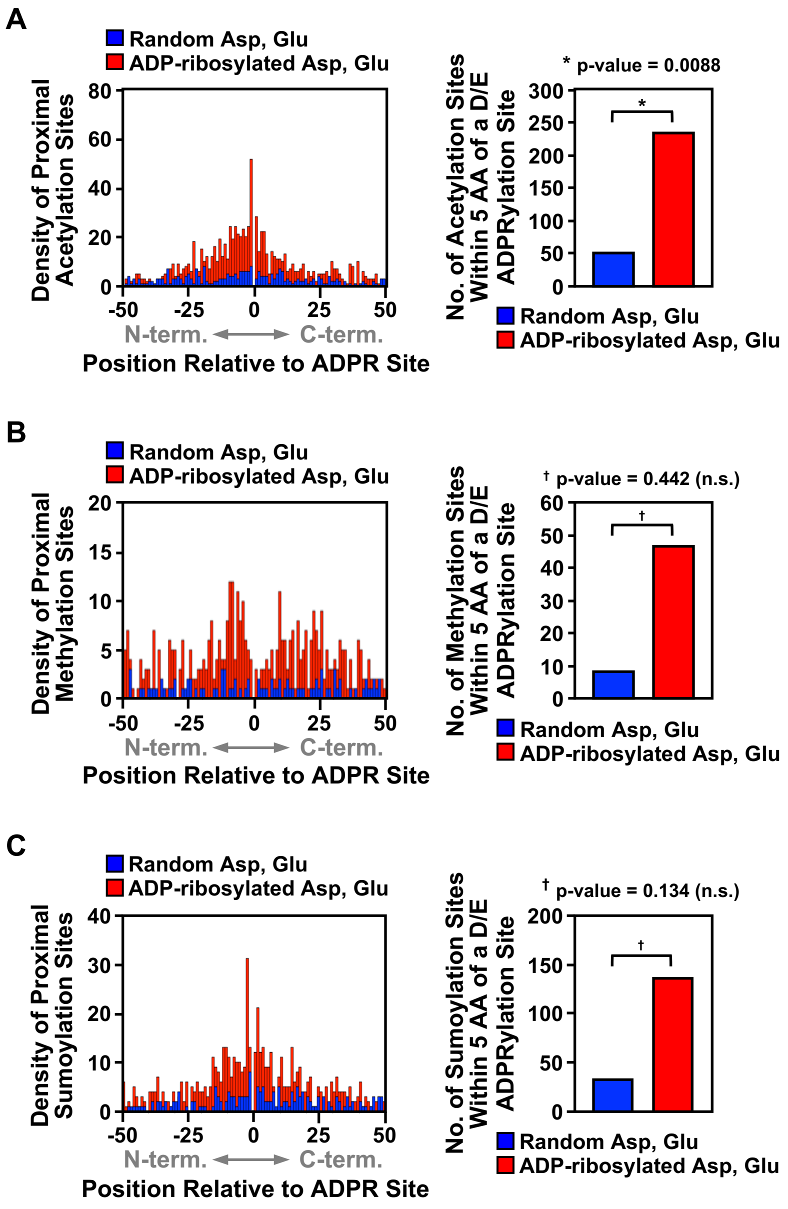

**Figure S4. Incidence of PTMs relative to ADPRylation sites.**
**(A - C)** Histograms (*left panels*) of the 2D relationship between the nearest incidence of known acetylation (A), methylation (B), or sumoylation (C) sites within ± 50 amino acids from the ADPRylation sites (Asp and Glu) identified herein. A random set of Asp and Glu residues were used as control. Bar graphs (*right panels*) showing quantification of each PTM within 5 amino acids of the ADPRylated site. A random set of Asp and Glu residues were used as control. Fisher's exact test was utilized to examine differences and p-values are noted.
*[Related to Figure 3A and 3B]*
